# Predicting coordinates of peptide features in raw timsTOF data with machine learning for targeted extraction reduces missing values in label-free DDA LC-MS/MS proteomics experiments

**DOI:** 10.1101/2022.04.25.489464

**Authors:** Daryl Wilding-McBride, Giuseppe Infusini, Andrew I. Webb

## Abstract

1

The determination of relative protein abundance in label-free data dependant acquisition (DDA) LC-MS/MS proteomics experiments is hindered by the stochastic nature of peptide detection and identification. Peptides with an abundance near the limit of detection are particularly effected. The possible causes of missing values are numerous, including; sample preparation, variation in sample composition and the corresponding matrix effects, instrument and analysis software settings, instrument and LC variability, and the tolerances used for database searching.

There have been many approaches proposed to computationally address the missing values problem, predominantly based on transferring identifications from one run to another by data realignment, as in MaxQuant’s matching between runs (MBR) method, and/or statistical imputation. Imputation transfers identifications by statistical estimation of the likelihood the peptide is present based on its presence in other technical replicates but without probing the raw data for evidence.

Here we present a targeted extraction approach to resolving missing values without modifying or realigning the raw data. Our method, which forms part of an end-to-end timsTOF processing pipeline we developed called Targeted Feature Detection and Extraction (TFD/E), predicts the coordinates of peptides using machine learning models that learn the delta of each peptide’s coordinates from a reference library. The models learn the variability of a peptide’s location in 3D space from the variability of known peptide locations around it. Rather than realigning or altering the raw data, we create a run-specific ‘lens’ through which to observe the data, targeting a location for each peptide of interest and extracting it. By also creating a method for extracting decoys, we can estimate the false discovery rate (FDR). Our method outperforms MaxQuant and MSFragger by achieving substantially fewer missing values across an experiment of technical replicates. The software has been developed in Python using Numpy and Pandas and open sourced with an MIT license (DOI 10.5281/zenodo.6513126) to provide the opportunity for further improvement and experimentation by the community. Data are available via ProteomeXchange with identifier PXD030706.

**Author Summary:** Missed identifications of peptides in data-dependent acquisition (DDA) proteomics experiments are an obstacle to the precise determination of which proteins are present in a sample and their relative abundance. Efforts to address the problem in popular analysis workflows include realigning the raw data to transfer a peptide identification from one run to another. Another approach is statistically analysing peptide identifications across an experiment to impute peptide identifications in runs in which they were missing.

We propose a targeted extraction technique that uses machine learning models to construct a run-specific lens through which to examine the raw data and predict the coordinates of a peptide in a run. The models are trained on differences between observations of confidently identified peptides in a run and a reference library of peptide observations collated from multiple experiments. To minimise the risk of drawing unsound experimental conclusions based on an unknown rate of false discoveries, our method provides a mechanism for estimating the false discovery rate (FDR) based on the misclassification of decoys as target features. Our approach outperforms the popular analysis tool suites MaxQuant and MSFragger/IonQuant, and we believe it will be a valuable contribution to the proteomics toolbox for protein quantification.

## 3 Introduction

A common challenge with DDA analysis of LC-MS/MS proteomics experiments is missing values (1). A value is considered missing in an experiment with technical replicates if a peptide was not identified in a run, but it was identified in one or more of the other runs. A missing value can occur for one of two reasons. It may be missing at random (MAR) for causes that are unrelated to its abundance, such as instrument fluctuations, LC variability, or a misidentification of the feature (2). Alternatively, it may be missing not at random (MNAR) because its abundance lies near the detection limit of the instrument (3). Popular solutions to missing values include data realignment to transfer identifications between runs, and statistical imputation.

Examples of transferring identifications between runs in commonly used free-to-download tools for processing timsTOF data can be found in MaxQuant (4) and MSFragger/IonQuant (5). Their approach is to match peptide features between runs through an alignment of the raw data. MaxQuant MBR applies windows for each identification in m/z, retention time, and mobility and find matching features. The m/z window is defined by the mass tolerances determined during mass recalibration; a wider window is used for retention time (15 minutes by default) and mobility. The distribution of deltas between runs is determined and a nonlinear calibration is applied to retention time and mobility, followed by feature matching with low tolerance in three dimensions (6). Thus, MaxQuant MBR will transfer an identified peptide from a donor run to an acceptor run if an unidentified feature in the acceptor run can be found with the same charge state in a specified window in m/z, retention time, and mobility, even if the acceptor run had no associated fragment ion information for that feature (7).

Without the assurance of an identification using fragment ions, false transfers can be an issue. Lim et al designed an experiment with two samples of Human cell lysate and yeast cell lysate to determine how often MaxQuant’s MBR algorithm incorrectly transfers an identification from one run to another. They analysed each sample with and without MBR enabled, and found MBR can introduce up to 44% false transfers, though only 2.7% of those remained after processing by MaxQuant’s LFQ analysis (8).

MSFragger’s transfer of identifications between runs is done by its companion tool IonQuant. As a measure to address the false positives of MaxQuant MBR approach, IonQuant’s MBR takes an approach whereby the FDR can be controlled. IonQuant calculates the correlation between runs by examining the amount of ion overlap in retention time, intensity, and mobility (9). Target ranges in each dimension for each ion whose identification is to be transferred are determined by calculating the median absolute deviation of the list of donor run ions. The FDR is estimated by creating decoy ions with the same retention time and mobility but shifted m/z. For the transferred and decoy ions, quality scores are calculated based on isotopic distribution in reference to the theoretical. Using a training set comprising target features identified using fragment spectra and decoys created by targets shifted in m/z, a classifier is trained and then used to classify transferred ions.

In the 3D space of timsTOF data, the position of a peptide feature will slightly vary from run to run in m/z, mobility, and retention time. This is due to electronic variation in the instrument with ambient temperature and humidity, variations in the mechanics and the chemistry of the liquid chromatography, and the interaction of peptide molecules in the ion optics (10). A peptide’s position in 3D space in a new run, however, is predictable, based on the variations we can observe in high abundance, confidently identified peptides over many runs.

As part of an analysis workflow named Targeted Feature Detection and Extraction (TFD/E), we developed an alternative to MBR that uses targeted extraction of a peptide at coordinates estimated by machine learning models trained on a library of prior peptide observations (Figure 1). An important point of differentiation between this approach and MBR is targeted extraction does not modify the raw data. Like a run-specific ‘lens’ into the raw data, the coordinate estimation models learn the run-specific deviations to accurately estimate the coordinates of each peptide in three dimensions (m/z, retention time, and mobility). We believe that by avoiding the alignment of data performed in other approaches our approach preserves the sanctity of the raw data and subtle multidimensional deviations that are more reliably taken into account by machine learning models than by hand-crafted predetermined tolerances. A target-decoy classifier learns how to distinguish between a good quality, identified feature and a false feature, and we use it to estimate the FDR and produce the final list of identified peptides in each run.

**Figure 1.**
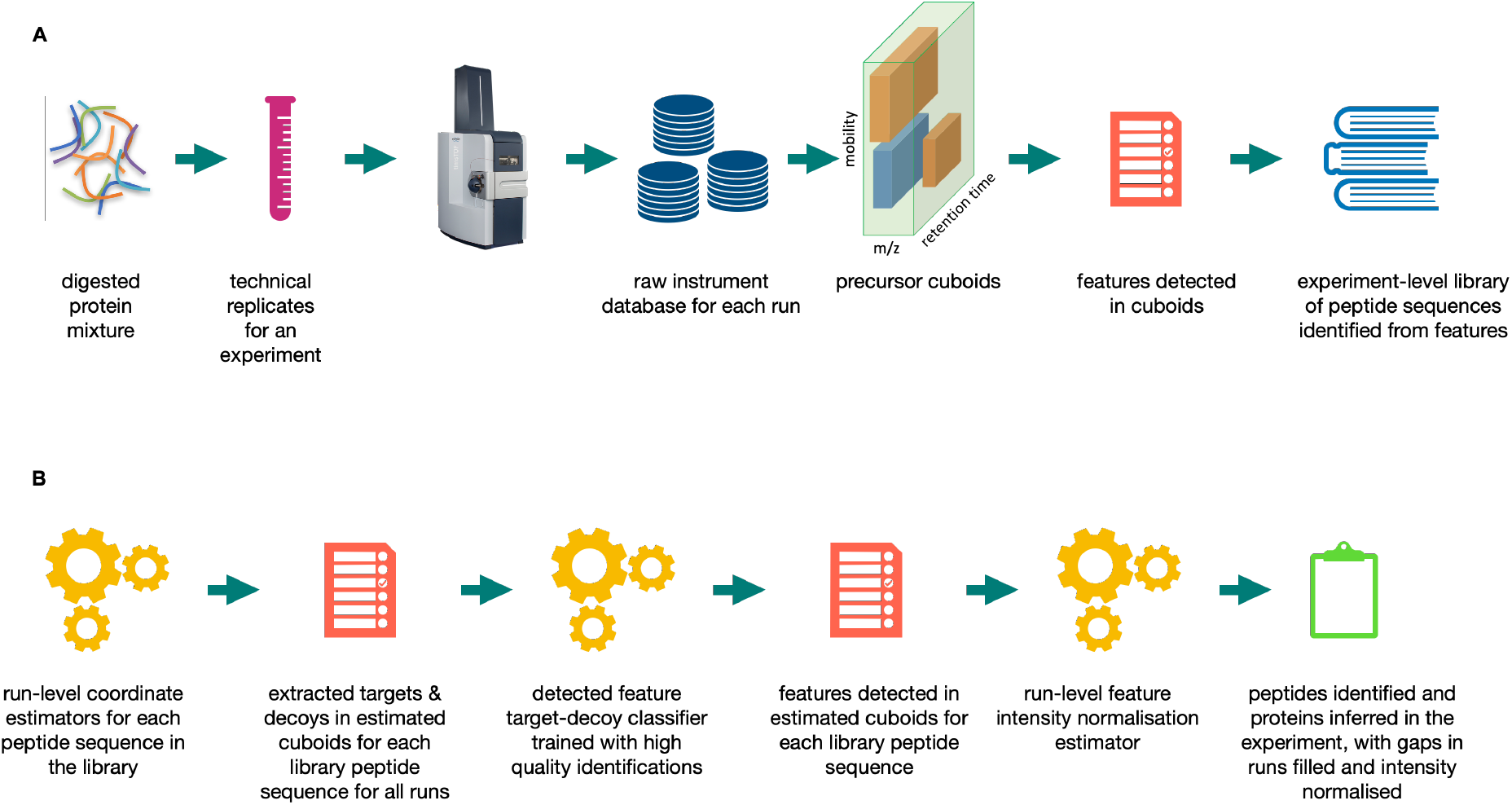
Schematic of the end-to-end TFD/E process flow. (A) The raw data from sample processing in the timsTOF is analysis with feature detection and identification to create an experiment-wide library of the peptides observed. (B) Coordinate estimator machine learning models predict the 3D location of each peptide in the library along with its decoy. A target-decoy classifier is used to estimate the FDR and determine the final list of peptide identifications for the experiment.

We show that using machine learning models to extract features from estimate coordinates with controlled FDR can achieve substantially fewer missing values than the MBR approaches of MaxQuant and MSFragger/IonQuant. We conclude that TFD/E offers a viable alternative to the currently popular approaches to addressing missing values in DDA analysis of LC-MS/MS proteomics experiments.

## 4 Results

### 4.1 Experimental design

To facilitate the comparison of peptide identifications and their intensity, we designed an experiment comprising technical replicates of proteome mixtures in different concentrations. A three-organism dataset was generated by mixing proteomes of *S. cerevisiae, H. sapiens*, and *E. coli*. The dataset comprised three experimental conditions with different protein ratios, each with ten technical replicates: 2:1:0 (*S. cerevisiae*), 1:1:1 (*H. sapiens*), and 1:4:0 (*E. coli*), named YHE211, YHE114, and YHE010 respectively. The dataset was analysed on a Bruker timsTOF Pro.

### 4.2 Building the peptide library

We use observations of peptides identified in at least one run to determine the experiment-wide mean and standard deviation of their position in m/z, retention time, and mobility to build a database of peptides for an experiment (Figure 1A). We also collate information about the base width of the peptides’ monoisotopic peaks in the retention time and mobility dimensions (Table 1).

**Table 1.**
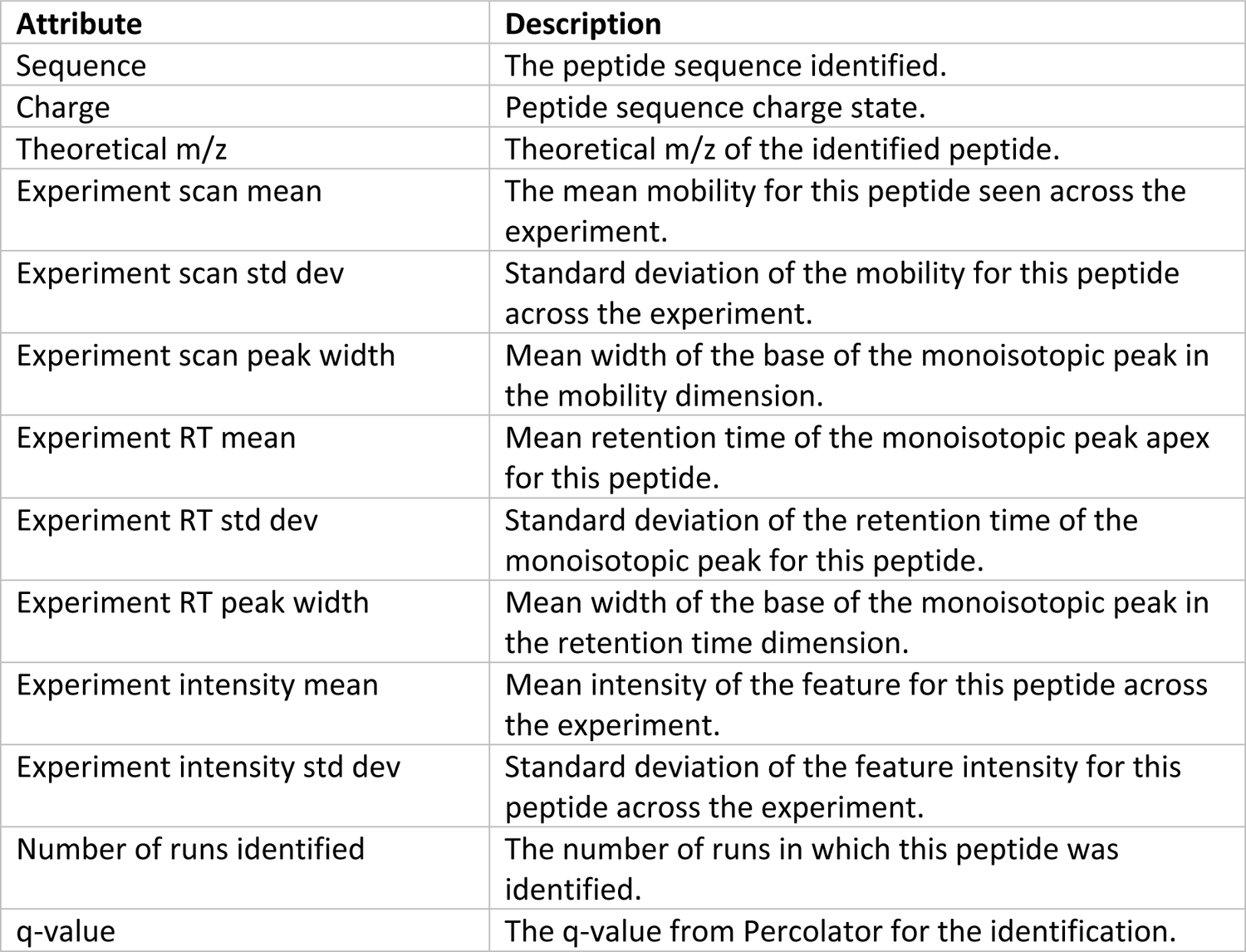
Attributes for each sequence collected in the peptide library

### 4.3 Building the coordinate estimators

Three Gradient Boosted Regressors (11) are trained to estimate feature coordinates in each run, one model each for the m/z, mobility, and retention time dimensions, and a set of models for each run. The training set for the models is created by taking the peptide identifications from the sequence library if the peptide was identified in more than half the runs. For each model’s output variable, we use delta from the experiment mean rather than absolute values to facilitate model convergence. The input and output variables for the models are shown in Table 2.

**Table 2.** Input variables (X) and output variable (y) for each coordinate estimator model.

| X | y |  |  |
| --- | --- | --- | --- |
|  | m/z | Mobility | Retention time |
| Theoretical m/z<br>Experiment RT mean<br>Experiment RT standard deviation<br>Experiment scan mean<br>Experiment scan standard deviation | Delta m/z (ppm) | Delta scan | Delta retention time (seconds) |
| ▪ Experiment intensity mean<br>▪ Experiment intensity standard deviation |  |  |  |

For an example run in the YHE211 experiment (P3856_YHE211_1), there were 11,016 unique peptides with a q-value lower than 0.01. Of these peptides, 8,955 were identified in more than half the technical replicates. To maximise the diversity of examples to be used for training, 98% of these were used for the training set, and 2% of these were set aside for the test set. The coordinate estimates from these models for this run against the test set achieved a mean absolute error (i.e., delta from the experiment mean) of 0.7664 m/z ppm, 0.0094 scan, and 0.0008 seconds retention time.

### 4.4 Using the coordinate estimators to extract features

The coordinate estimators are used to estimate the peptide coordinates - monoisotopic m/z, apex in retention time, and apex in mobility - in each run. The peptide’s charge state is known, as well as the experiment-wide mean width at the base of its monoisotopic peak in retention time and mobility. Allowing for up to seven isotopes for the purpose of defining the bounds of a feature search cuboid, a region of raw data is formed around the estimated coordinates in which to find the target feature.

Raw data points belonging to the monoisotope are grouped by taking 5 ppm either side of the estimated monoisotopic m/z. The intensity-weighted m/z centroid is used as the extracted m/z of the peak.

Points belonging to the feature’s isotopes are gathered from expected isotope spacing from the monoisotopic m/z according to the peptide’s charge state:

~~~
expected_spacing_mz = carbon_mass_difference / charge
~~~

where `carbon_mass_difference = 1.003355`, the mass difference between Carbon-12 and Carbon-13 isotopes, in Da.

Having gathered the raw points belonging to each isotopic peak, we find the apex of the points for the monoisotopic peak in retention time and mobility. In mobility we do this by summing the points occurring on the same scan, smoothing the points with a Savitzky-Golay filter (12), and find the peaks and troughs with the PeakUtils Python library (13). We repeat the same steps to find the apex of the peak in retention time, instead summing the peak’s raw points that occur in the same frame.

Having extracted the target feature and its attributes from the raw data at the estimated coordinates, feature metrics (listed in Table 5) are calculated to be used for the target-decoy classifier input variables.

### 4.5 Extracting decoys

In noisy raw data, it is feasible to start with a random coordinate and extract points that could fit the pattern of a monoisotopic peak followed by a series of isotopic peaks at the correct spacing in m/z for the feature’s charge state. To estimate the false discovery rate

(FDR) of the feature extraction, a target-decoy classifier is used. The rationale for creating decoys in close proximity to targets is to include in the FDR calculation the possibility of a “near miss” of the target, where a false peptide feature in similar local noise conditions will be extracted from a location nearby. To generate metrics for decoys, decoy coordinates are calculated by offsetting the target coordinates by a random amount within predetermined limits in each dimension, to calculate decoy coordinates that are in the neighbourhood of a real feature, but sufficiently offset from it (Table 3).

**Table 3.** Offsets applied to the target coordinate to determine the decoy coordinate

| **Decoy coordinate** | **Offset from the target coordinate** |
| --- | --- |
| Monoisotopic m/z | Random delta between +/- 20 ppm |
| Apex of the monoisotopic peak in the mobility dimension | Random delta between +/- 2 peak widths, where peak width is the library-wide mean width in the mobility dimension for that peptide. |
| Apex of the monoisotopic peak in the retention time dimension | Random delta between +/- 2 peak widths, where peak width is the library-wide mean width in the retention time dimension for that peptide. |

For each peptide in the library, metrics are calculated for the false feature extracted from the decoy coordinates.

### 4.6 Building a target-decoy classifier for extracted features

The training set for the target-decoy classifier is composed of the metrics calculated for the target and decoy of each peptide in the library that was identified in more than half the technical replicates. A Gradient Boosted Classifier (14) is used for the model. As the number of targets with a full set of extractable metrics is expected to exceed the number of extractable decoys, the targets are downsampled at random to balance the training set. 10% of the set is set aside for the test set, and 90% are used for training. In a typical analysis of the YHE211 technical replicates, the classifier has an FDR of 1.2% (Table 4).

**Table 4.** Classification report for the target-decoy classifier.

|  | **Precision** | **Recall** | **F1 Score** | **Support** |
| --- | --- | --- | --- | --- |
| **Target** | 0.98 | 0.99 | 0.98 | 8859 |
| **Decoy** | 0.99 | 0.98 | 0.98 | 8590 |

### 4.7 Classifying the extracted features as target or decoy

The target-decoy classifier is used to classify the target metrics from each run in the experiment for each sequence in the peptide library. Each unique occurrence of a peptide in each run from the first pass of identifications and then by targeted extraction were counted for the YHE211 and YHE114 technical replicates (Figure 2A). Compared with the first pass of identification in each run, targeted extraction increases the yield of peptides by an average of 58% across the experiment.

**Figure 2.**
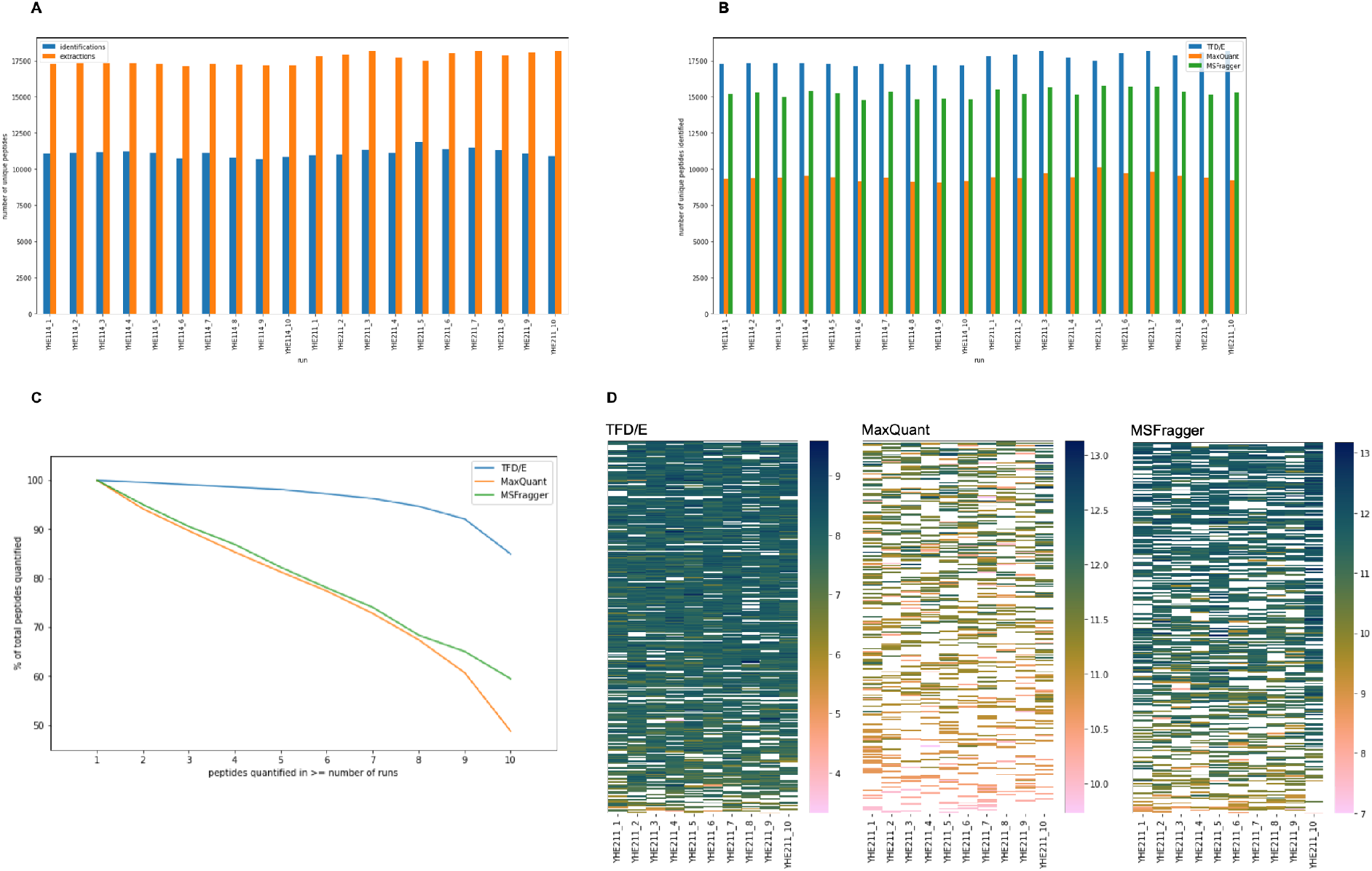
(A) The number of unique peptide identifications and extractions in each run of the YHE211 and YHE114 experiments, (B) Comparing the unique peptides identified in each run by TFD/E, MaxQuant, and MSFragger, (C) The percentage of peptides without missing values as a function of run numbers for the experiment condition YHE211, (D) Lowest 500 peptides by mean intensity for YHE211 from TFD/E, MaxQuant, and MSFragger. Blank cells represent missing values.

To compare with the identifications with other commonly used tool suites, we analysed the same technical replicates with MaxQuant (Version 1.6.17) and MSFragger (FragPipe 13.0, MSFragger version 3.0, Philosopher version 3.2.9), each with MBR enabled. TFD/E yields 86% more peptides per run compared to MaxQuant, and 15% per run more than MSFragger (Figure 2B).

As well as a higher yield of unique peptides in each run, TFD/E has fewer missing values across the experiment. The percentage of the total peptides quantified in more than N runs of the YHE211 samples by TFD/E, MaxQuant, and MSFragger was plotted as a function of the number of runs in which the peptides were extracted (Figure 2C). While MaxQuant and MSFragger have a steady decrease in percentage of peptides quantified as the number of runs increases, TFD/E sustains a higher percentage of peptides quantified as the number of runs increases, only dropping below 90% of the total number of peptides for 9 runs or more.

For the comparison of false discovery and false extraction/transfer, the following definitions were used:

• The “false discovery rate” (FDR) is the number of unique non-human peptides as the proportion of all unique peptides identified. A unique peptide is a unique combination of modified sequence and charge state.

• A “transfer” is a successful targeted extraction from a run in which the peptide was not detected and identified.

• “All possible false transfers” is the total number of runs a falsely-identified peptide could have been transferred. So if a non-human peptide was detected and identified in 2 runs, the number of possible false transfers was 10-2=8.

To examine the FDR of each approach, the human-only YHE010 technical replicates were analysed by TFD/E, MaxQuant, and MSFragger using the same FASTA database containing Yeast, HeLa, and *E*.*coli* proteins. The FDR for TFD/E, MaxQuant, and MSFragger was determined to be 1.3%, 0.3%, and 0.5% respectively. As TFD/E uses an FDR cutoff of 1% for identifications, its result is consistent with what one would expect.

Once a false peptide is included in the peptide library for extraction, the false extraction/transfer rate as a proportion of all possible false transfers was 81% for TFD/E, and 39% for MaxQuant. The false transfer rate for MSFragger was not determined as it does not report the distinction between an identification and a transfer.

From this we conclude that TFD/E is better than MaxQuant at extracting/transferring a detected feature, even when that feature is not correctly identified. This means the FDR in TFD/E has an amplifying effect on reducing missing values, filling gaps for a false identification.

To compare how each tool suite performs for very low abundance peptides, the 500 lowest intensity peptides extracted by each tool for the YHE211 samples were plotted by run (Figure 2D). TFD/E has the fewest missing values for this group of peptides, highlighting its ability to consistently extract peptides even of very low abundance.

The coefficient of variance for peptide intensity across the YHE211 runs was calculated for TFD/E, MaxQuant, and MSFragger (Figure 3A), and we found that TFD/E has similar performance.

**Figure 3.**
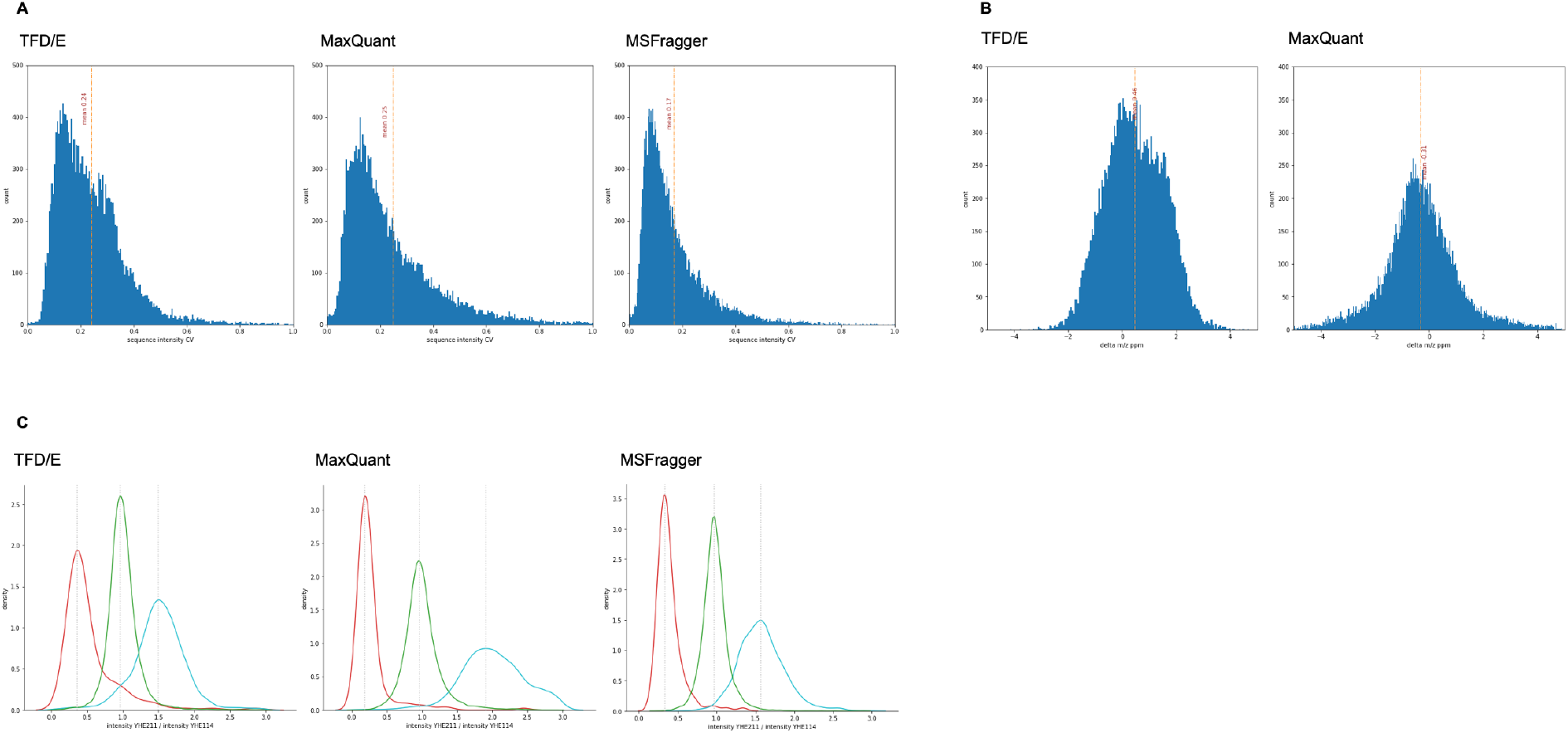
(A) Peptide intensity coefficient of variance for condition YHE211 processes by TFD/E, MaxQuant, and MSFragger. (B) Peptide mean delta mass for the experiment condition YHE211 for TFD/E and MaxQuant. (C) Density plot of Human (green), yeast (blue), and E.coli (red) peptide intensity ratios between experiment conditions YHE211 and YHE114 for TFD/E, MaxQuant, and MSFragger.

The delta mass ppm between extractions and the theoretical (Figure 3B) is slightly higher for TFD/E than MaxQuant but still well within +/- 2 ppm. MSFragger delta mass from the theoretical is not shown because it is not reported for identifications transferred between runs with MBR.

The ratios of the mean intensity of each peptide detected in both YHE211 and YHE114 experiment conditions were compared (Figure 3C). The intensity ratios of peptides from MaxQuant are the closer to the expected than TFD/E. The small bumps around 1.0 in the E.coli (red) and Yeast (blue) traces of the TFD/E density plot suggest that some Human peptides have been misidentified. This could be caused by peptide feature false discovery or by mis-extraction (e.g., extracting a proximate human peptide of the same charge state from the vicinity of the estimated m/z, retention time, and mobility for a targeted E. coli peptide).

## 5 Discussion

In this work we have shown that our method for targeted extraction substantially decreases missing values in technical replicates compared to MaxQuant and MSFragger with MBR enabled. An advantage of our approach over MBR techniques is it constructs a run-specific lens into the data without altering it, and it does not rely on transferring an identification from a donor run to an acceptor run. By extracting decoys as well as targets we can estimate the false discovery rate.

Another appealing aspect to this approach is that the peptide library from which the coordinate estimators are trained can be constructed by several means and not just by taking the mean of observations of technical replicates within a single experiment. A peptide library could be constructed from a peptide knowledge base collated from a broad range of experiments, or from predicted peptide attributes. Using the peptide library to train the coordinate estimators and extract peptides is completely independent of how the library was created. The further application of this is an interesting area for additional research. While this work has focused on predicting the coordinates of precursor ions, the coordinates of fragment ions as deltas from a reference peptide library may also be predicted, opening the possibility the approach could be applied to DIA extraction. We believe that TFD/E is an attractive alternative to common approaches to the missing values problem, and thereby it promises to be a valuable addition to the proteomics toolbox for improved protein quantification in LC-MS/MS experiments.

## 6 Materials and Methods

### 6.1 Sample preparation

Commercial tryptic digests of *S*.*cerevisiae* (Yeast, Promega, #V746A), human K562 cells (Promega, #V695A) and *E*.*coli* (MassPREP standard, Waters, #186003196) were reconstituted in 2% ACN/0.1% FA to final concentration of 0.1 μg/ul. To generate the hybrid proteome samples, purified peptides from each of the three species were combined in different proportions as previously described (40) and as follows: sample YHE211 consisted of 30% w/w Yeast (3 μg), 65% w/w Human (6.5 μg) and 5% w/w *E.coli* (0.5 μg); sample YHE114 consisted of 15% w/w Yeast (1.5 μg), 65% w/w Human (6.5 μg) and 20% w/w *E.coli* (2 μg); sample YHE010 consisted of 0% w/w Yeast (0 μg), 100% w/w Human (6.5 μg), and 0% w/w *E.coli* (0 μg). Ten replicates of each proteome mixture were subjected to LC-MS/MS analysis on a timsTOF Pro mass spectrometer.

### 6.2 LC-MS methods

The digested proteome mixtures were separated by nanoflow reverse-phase chromatography (i.d. 75 μm, o.d. 360 μm × 25 cm length, 1.6 μm C18 beads - IonOpticks, Australia) using a nanoflow HPLC (M-class, Waters). The HPLC was coupled to a timsTOF Pro mass spectrometer (Bruker Daltonics, Bremen) using a CaptiveSpray source. Peptides were loaded directly onto the column at a constant flow rate of 400 nL/min with buffer A (99.9% Milli-Q water, 0.1% FA) and eluted with a 20-minute linear gradient from 2% to 34% buffer B (99.9% ACN, 0.1% FA).

The timsTOF Pro was operated in PASEF mode using Compass Hystar 5.1 and otofControl settings were as follows: Mass Range 100 to 1700m/z, 1/K0 Start 0.85 V·s/cm^2^ End 1.3 V·s/cm^2^, Ramp time 100 ms, Lock Duty Cycle to 100%, Capillary Voltage 1600V, Dry Gas 3 l/min, Dry Temp 180°C, PASEF settings: 4 MS/MS scans (total cycle time 1.27sec), charge range 0-5, active exclusion for 0.4 min, Scheduling Target intensity 24000, Intensity threshold 2500.

### 6.3 Mass Spectrometry Analysis

The raw data was extracted from the instrument database and processed with our bespoke software for feature detection. For peptide identification and protein inference we used Crux Comet and Percolator. The mass tolerance for the initial search was 20 ppm; the search following mass recalibration was 4.5 ppm. Settings were a maximum of 2 missed cleavages, a bin tolerance of 0.02 Da for fragment ions, a bin offset of 0, and a default peak shape. The MH+ peptide mass range for analysis was 700-5000 Da.

The FASTA database was created for the Yeast/HeLa/E.coli proteome mixtures by combining databases for each proteome from UniProtKB (16–18).

### 6.4 Software

The software was written in Python 3.8. The key libraries used were Pandas 1.3.1 for data filtering and interface file input/output, sklearn 1.0 for the Gradient Boosted Regressors and Classifier, and Ray 1.5.2 for parallel processing. Algorithm prototyping was done in Jupyter notebooks (jupyter-core 4.6.3).

Software validation work was performed on a PC with a 12-core Intel i7 6850K processor and 64 GB of memory running Ubuntu 20.04.

Readers are encouraged to browse the source code in the GitHub repository (DOI 10.5281/zenodo.6513126) for a detailed understanding of the algorithms and implementation approach. The Jupyter notebooks developed to generate the figures in this paper are also available in the repository.

## 7. Acknowledgments

The authors gratefully acknowledge the contributions of the following:

• Laura Dagley and Sukhdeep Kaur Spall at WEHI, who prepared the samples for the experiment and processed them with the timsTOF.

• Sven Brehmer at Bruker for discussions and guidance about processing timsTOF data.

## 9 Appendices

**Table 5.**
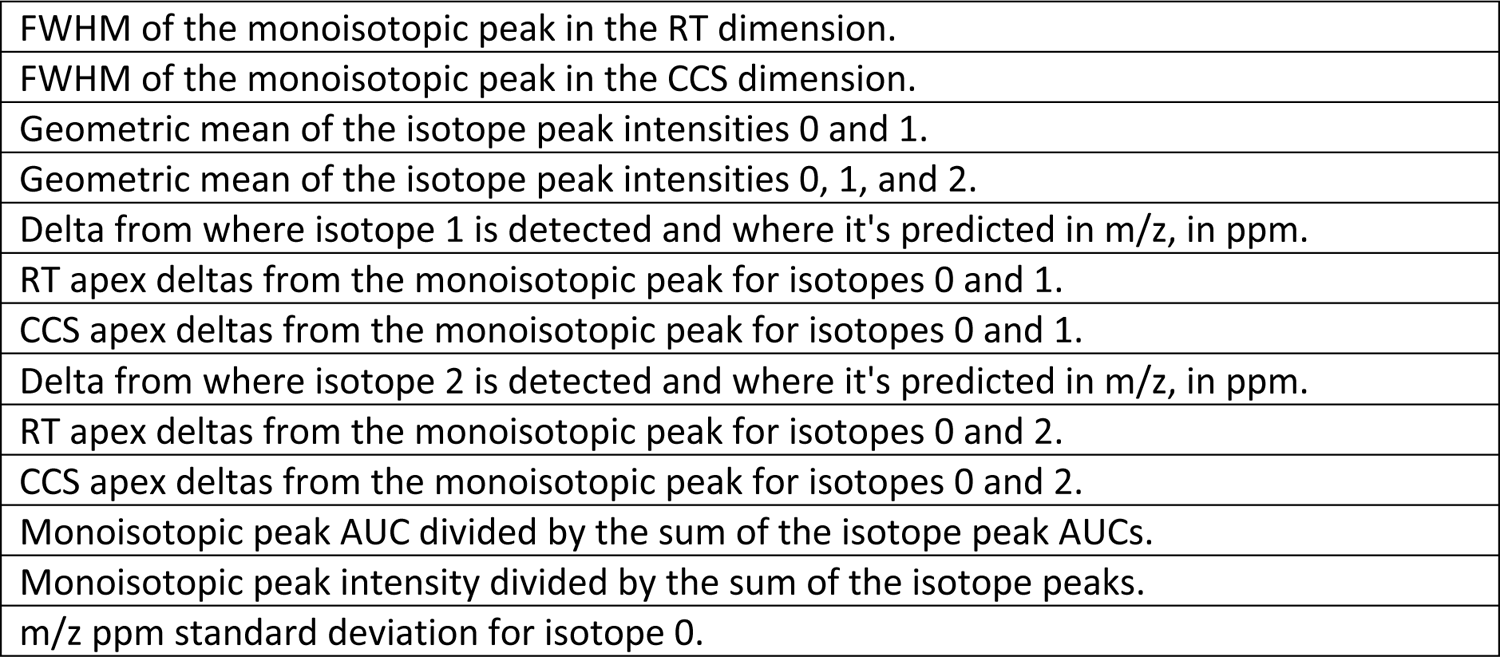
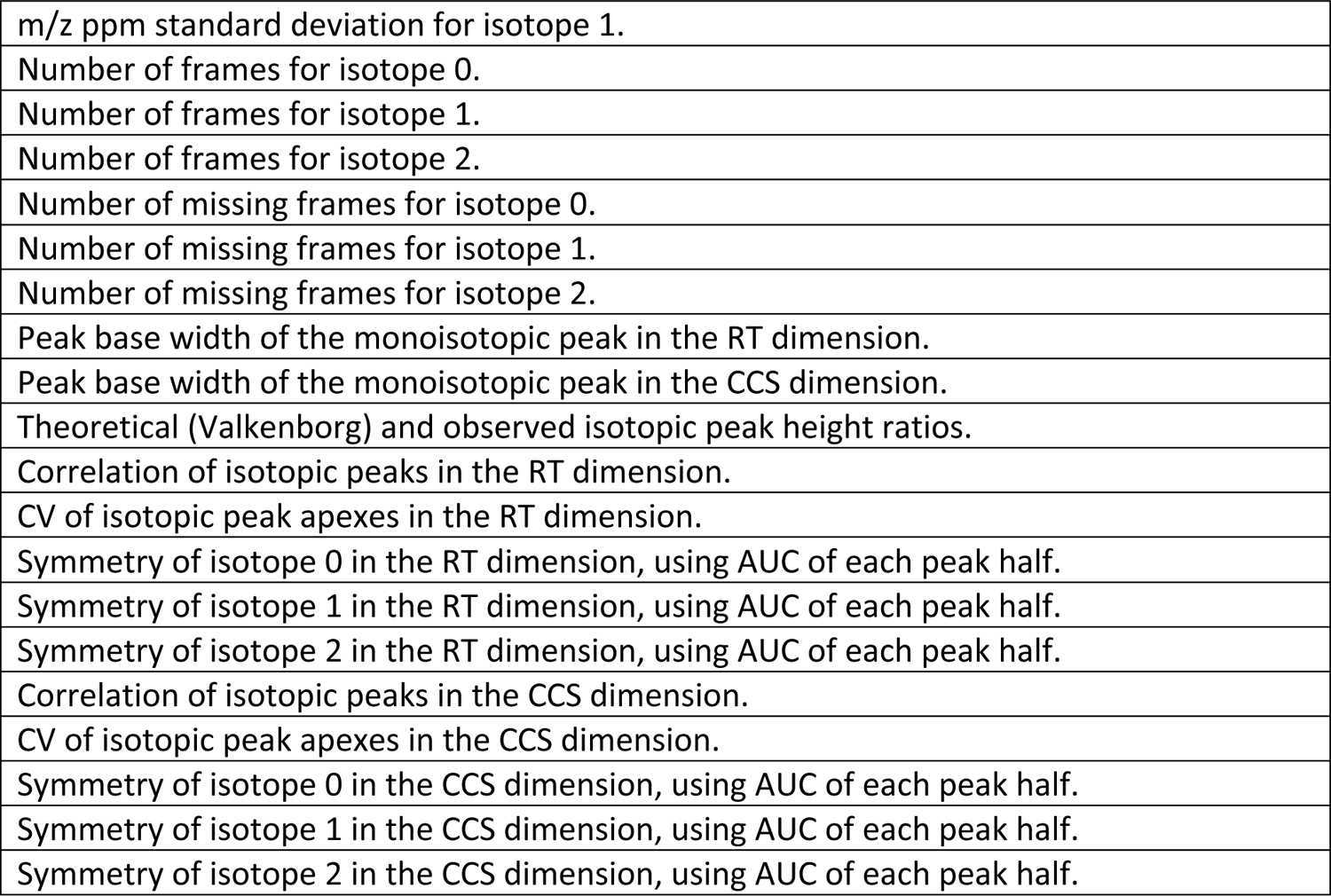
Feature metrics extracted for the target-decoy classifier

|  |
| --- |
| FWHM of the monoisotopic peak in the RT dimension. |
| FWHM of the monoisotopic peak in the CCS dimension. |
| Geometric mean of the isotope peak intensities 0 and 1. |
| Geometric mean of the isotope peak intensities 0, 1, and 2. |
| Delta from where isotope 1 is detected and where it's predicted in m/z, in ppm. |
| RT apex deltas from the monoisotopic peak for isotopes 0 and 1. |
| CCS apex deltas from the monoisotopic peak for isotopes 0 and 1. |
| Delta from where isotope 2 is detected and where it's predicted in m/z, in ppm. |
| RT apex deltas from the monoisotopic peak for isotopes 0 and 2. |
| CCS apex deltas from the monoisotopic peak for isotopes 0 and 2. |
| Monoisotopic peak AUC divided by the sum of the isotope peak AUCs. |
| Monoisotopic peak intensity divided by the sum of the isotope peaks. |
| m/z ppm standard deviation for isotope 0. |
| m/z ppm standard deviation for isotope 1. |
| Number of frames for isotope 0. |
| Number of frames for isotope 1. |
| Number of frames for isotope 2. |
| Number of missing frames for isotope 0. |
| Number of missing frames for isotope 1. |
| Number of missing frames for isotope 2. |
| Peak base width of the monoisotopic peak in the RT dimension. |
| Peak base width of the monoisotopic peak in the CCS dimension. |
| Theoretical (Valkenborg) and observed isotopic peak height ratios. |
| Correlation of isotopic peaks in the RT dimension. |
| CV of isotopic peak apexes in the RT dimension. |
| Symmetry of isotope 0 in the RT dimension, using AUC of each peak half. |
| Symmetry of isotope 1 in the RT dimension, using AUC of each peak half. |
| Symmetry of isotope 2 in the RT dimension, using AUC of each peak half. |
| Correlation of isotopic peaks in the CCS dimension. |
| CV of isotopic peak apexes in the CCS dimension. |
| Symmetry of isotope 0 in the CCS dimension, using AUC of each peak half. |
| Symmetry of isotope 1 in the CCS dimension, using AUC of each peak half. |
| Symmetry of isotope 2 in the CCS dimension, using AUC of each peak half. |

